# Similar evolutionary trajectories for retrotransposon accumulation in mammals

**DOI:** 10.1101/091652

**Authors:** Reuben M Buckley, R Daniel Kortschak, Joy M Raison, David L Adelson

**Affiliations:** Department of Genetics and Evolution, The University of Adelaide, North Tce, 5005, Adelaide, Australia

**Keywords:** Transposable element, Genome Evolution, Genome Architecture

## Abstract

The factors guiding retrotransposon insertion site preference are not well understood. Different types of retrotransposons share common replication machinery and yet occupy distinct genomic domains. Autonomous long interspersed elements accumulate in gene-poor domains and their non-autonomous short interspersed elements accumulate in gene-rich domains. To determine genomic factors that contribute to this discrepancy we analysed the distribution of retrotransposons within the framework of chromosomal domains and regulatory elements. Using comparative genomics, we identified large-scale conserved patterns of retrotransposon accumulation across several mammalian genomes. Importantly, retrotransposons that were active after our sample-species diverged accumulated in orthologous regions. This suggested a similar evolutionary interaction between retrotransposon activity and conserved genome architecture across our species. In addition, we found that retrotransposons accumulated at regulatory element boundaries in open chromatin, where accumulation of particular retrotransposon types depended on insertion size and local regulatory element density. From our results, we propose a model where density and distribution of genes and regulatory elements canalise retrotransposon accumulation. Through conservation of synteny, gene regulation and nuclear organisation, mammalian genomes with dissimilar retrotransposons follow similar evolutionary trajectories.

## Introduction

An understanding of the dynamics of evolutionary changes in mammalian genomes is critical for understanding the diversity of mammalian biology. Most work on mammalian molecular evolution is on protein coding genes, based on the assumed centrality of their roles and because of the lack of appropriate methods to identify the evolutionary conservation of apparently non-conserved, non-coding sequences. Consequently, this approach addresses only a tiny fraction (less than 2%) of a species’ genome, leaving significant gaps in our understanding of evolutionary processes (ENCODE Project Consortium 2012; Lander et al. 2001). In this report we describe how large scale positional conservation of non-coding, repetitive DNA sheds light on the possible conservation of mechanisms of genome evolution, particularly with respect to the acquisition of new DNA sequences.

Mammalian genomes are hierarchically organised into compositionally distinct hetero- or euchromatic large structural domains (Gibcus and Dekker 2013). These domains are largely composed of mobile self-replicating non-long terminal repeat (non-LTR) retrotransposons; with Long INterspersed Elements (LINEs) in heterochromatic regions and Short INterspersed Elements (SINEs) in euchromatic regions (Medstrand et al. 2002). The predominant LINE in most mammals is the ~6 kb long L1. In many mammal genomes, this autonomously replicating element is responsible for the mobilisation of an associated non-autonomous SINE, usually ~300 bp long. Together, LINEs and SINEs occupy approximately 30% of the human genome (Lander et al. 2001), replicate via a well characterised RNA-mediated copy-and-paste mechanism (Cost et al. 2002) and co-evolve with host genomes (Kramerov and Vassetzky 2011; Chalopin et al. 2015; Furano et al. 2004).

The accumulation of L1s and their associated SINEs into distinct genomic regions depends on at least one of two factors. 1) Each element’s insertion preference for particular genomic regions and 2) the ability of particular genomic regions to tolerate insertions. According to the current retrotransposon accumulation model, both L1s and SINEs likely share the same insertion patterns constrained by local sequence composition. Therefore, their accumulation in distinct genomic regions is a result of region specific tolerance to insertions. Because L1s are believed to have a greater capacity than SINEs to disrupt gene regulatory structures, they are evolutionarily purged from gene-rich euchromatic domains at a higher rate than SINEs. Consequently, this selection asymmetry in euchromatic gene-rich regions causes L1s to become enriched in gene-poor heterochromatic domains (Lander et al. 2001; Graham and Boissinot 2006; Gasior et al. 2007; Kvikstad and Makova 2010).

An important genomic feature, not explored in the accumulation model, is the chromatin structure that surrounds potential retrotransposon insertion sites. Retrotransposons preferentially insert into open chromatin (Cost et al. 2001; Baillie et al. 2011), which is usually found overlapping gene regulatory elements. As disruption of regulatory elements can often be harmful, this creates a fundamental evolutionary conflict for retrotransposons; their immediate replication may be costly to the overall fitness of the genome in which they reside. Therefore, rather than local sequence composition or tolerance to insertion alone, retrotransposon accumulation is more likely to be constrained by an interaction between retrotransposon expression, openness of chromatin, susceptibility of a particular site to alter gene regulation, and the capacity of an insertion to impact on fitness.

To investigate the relationship between retrotransposon activity and genome evolution, we began by characterising the distribution and accumulation of non-LTR retrotransposons within placental mammalian genomes. Next, we compared retrotransposon accumulation patterns in eight separate evolutionary paths by ‘humanising’ the repeat content (see methods) of the chimpanzee, rhesus macaque, mouse, rabbit, dog, horse and cow genomes. Finally, we analysed human retrotransposon accumulation in large hetero- and euchromatic structural domains, focusing on regions surrounding genes, exons and regulatory elements. Our results suggest that accumulation of particular retrotransposon families follows from insertion into open chromatin found adjacent to regulatory elements and depends on local gene and regulatory element density. From this we propose a refined retrotransposon accumulation model in which random insertion of retrotransposons is primarily constrained by chromatin structure rather than local sequence composition.

## Materials and Methods

### Within species comparisons of retrotransposon genome distributions

Retrotransposon coordinates for each species were initially identified using RepeatMasker and obtained from either the RepeatMasker website or UCSC genome browser (Table S1) (Smit et al. 1996; Rosenbloom et al. 2015). We grouped retrotransposon elements based on repeat IDs used in Giordano *et al* (Giordano et al. 2007). Retrotransposon coordinates were extracted from hg19, mm9, panTro4, rheMac3, oruCun2, equCab2, susScr2, and canFam3 assemblies. Each species genome was segmented into 1 Mb regions and the density of each retrotransposon family for each segment was calculated. Retrotransposon density of a given genome segment is equal to a segments total number of retrotransposon nucleotides divided by that segments total number of mapped nucleotides (non-N nucleotides). From this, each species was organised into an *n*-by-*p* data matrix of *n* genomic segments and *p* retrotransposon families. Genome distributions of retrotransposons were then analysed using principle component analysis (PCA) and correlation analysis. For correlation analysis, we used our genome segments to calculate Pearson’s correlation coefficient between each pair-wise combination of retrotransposon families within a species.

### Across species comparisons of retrotransposon genome distributions

To compare genome distributions across species, we humanised a segmented query species genome using mapping coordinates extracted from net AXT alignment files located on the UCSC genome browser (Table S1). First, poorly represented regions were removed by filtering out genome segments that fell below a minimum mapping fraction threshold (Fig. 1a). Next, we used mapping coordinates to match fragments of query species segments to their corresponding human segments (Fig. 1b). From this, the retrotransposon content and PC scores of the matched query segments were humanised following equation 1 (Fig. 1c).

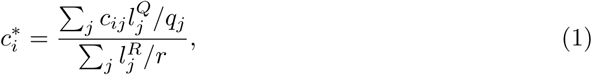

where *c_ij_* is the density of retrotransposon family *i* in query segment *j*, 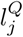 is the total length of the matched fragments between query segment *j* and the reference segment, 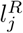 is the total length of the reference segment fragments that match query segment *j*, *q_j_* is the total length of the query segment *j*, and *r* is the total length of the reference segment. The result 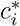 is the humanised coverage fraction of retrotransposon family *i* that can now be compared to a specific reference segment. Once genomes were humanised, Pearson’s correlation coefficient was used to determine the conservation between retrotransposon genomic distributions (Fig. 1d). Using the Kolmogorov-Smirnov test, we measured the effect of humanising by comparing the humanised query retrotransposon density distribution to the query filtered retrotransposon density distribution (Fig. 1e). The same was done to measure the effect of filtering by comparing the segmented human retrotransposon density distribution to the human filtered retrotransposon density distribution (Fig. 1f). Our Pearson’s correlation coefficients and P-values from measuring the effects of humanising and filtering were integrated into a heatmap (Fig. 1g). This entire process was repeated at different minimum mapping fraction thresholds to optimally represent each retrotransposon families genomic distribution in a humanised genome (fig S1).

**Figure 1.**
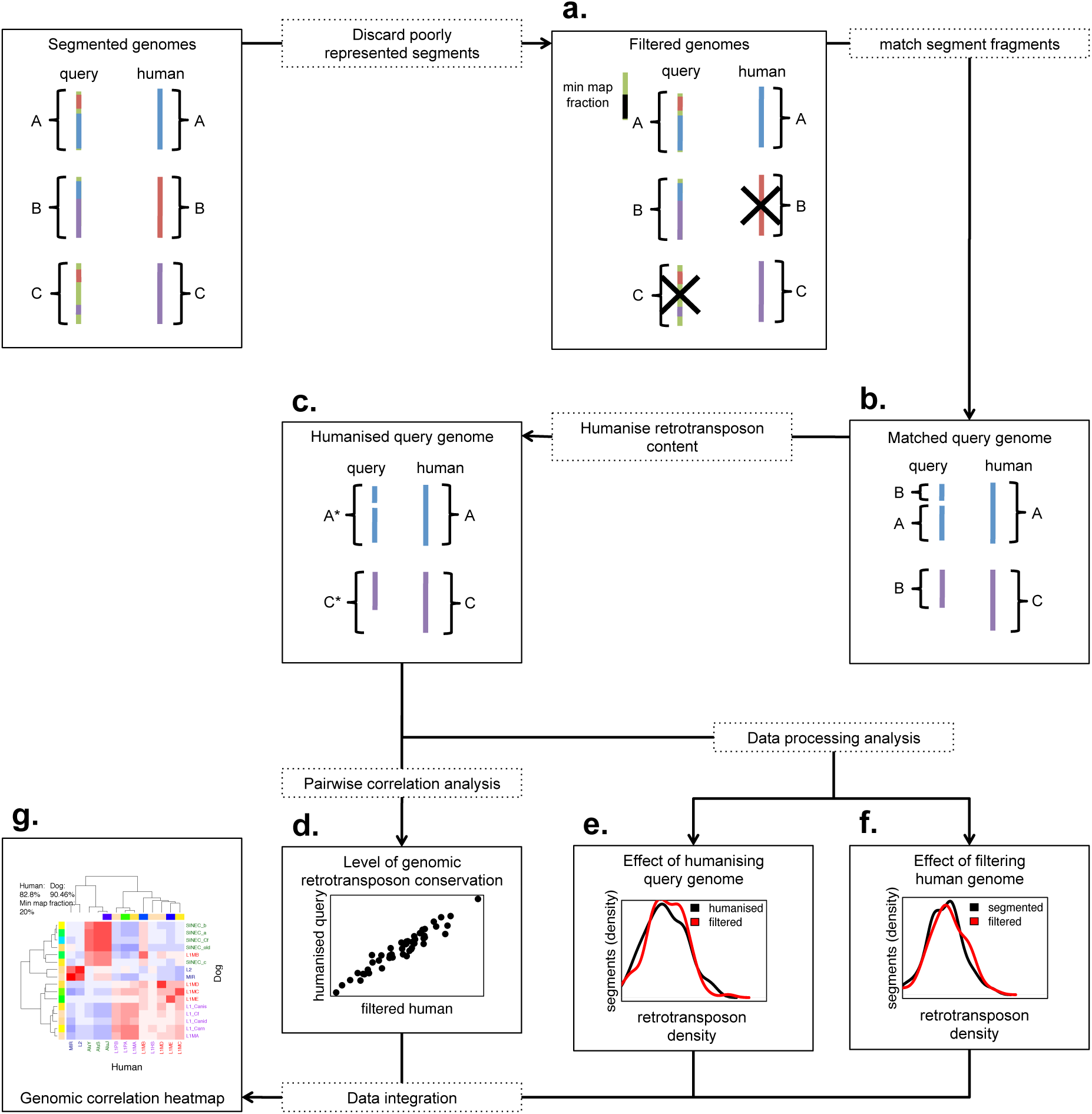
Overview of humanising retrotransposon distributions. **a**, Genomes are segmented and filtered according to a minimum mapping fraction threshold, removing poorly represented segments from both species. The black X shows which segments were not able to reach the minimum mapping fraction threshold. **b**, Fragments of query species’ genome segments are matched to their corresponding human genome segments using genome alignments. **c**, Query species genomes are humanised following equation 1. **d**, Pairwise genomic correlations are measured between each humanised retrotransposon family and each human retrotransposon family. **e**, The effect of humanising on retrotransposon density distributions is measured by performing a Kolmogorov-Smirnov test between the humanised query retrotransposon density distribution and the filtered query retrotransposon density distribution. **f**, The effect of filtering on retrotransposon density distributions is measured by performing a Kolmogorov-Smirnov test between the segmented human retrotransposon density distribution and the filtered human retrotransposon density distribution. **g**, The pairwise correlation analysis results and the P-values from the Kolmogorov-Smirnov tests are integrated into heatmaps (Fig. 4, S18-S22) that compare the genomic relationships of retrotransposons between species.

### Replication timing profiles, boundaries and constitutive domains

Genome-wide replication timing data for human and mouse were initially generated as part of the ENCODE project and were obtained from UCSC genome browser (Table S2-S3) (Yue et al. 2014; ENCODE Project Consortium 2012). For human genome-wide replication timing we used Repli-Seq smoothed wavelet signals generated by the UW ENCODE group (ENCODE Project Consortium 2012), in each cell-line we calculated the mean replication timing per 1Mb genome segment. For mouse genome-wide replication timing we used Repli-Chip wave signals generated by the FSU ENCODE group (Yue et al. 2014). Since two replicates were performed on each cell-line, we first calculated each cell-lines mean genome-wide replication timing and then used this value to calculate the mean replication timing per 1Mb genome segment. By calculating mean replication timing per 1 Mb segment we were able to easily compare large-scale genome-wide replication timing patterns across cell-lines. We obtained early replication domains (ERDs), late replication domains (LRDs) and timing transition regions (TTRs) from the gene expression omnibus (accession ID GSE53984) (Table S2). Replication domains for each dataset were identified using a deep neural network hidden Markov model (Liu et al. 2015). To determine RD boundary fluctuations of retrotransposon density, we defined ERD boundaries as the boundary of a TTR adjacent to an ERD. ERD boundaries from across each sample were pooled and retrotransposon density was calculated for 50 kb intervals from regions flanking each boundary 1 Mb upstream and downstream. Expected density and standard deviation for each retrotransposon group was derived from a background distribution generated by calculating the mean of 500 randomly sampled 50 kb genomic bins within 2000 kb of each ERD boundary, replicated 10000 times. To generate replication timing profiles for our ERD boundaries, we also calculated the mean replication timing per 50 kb intervals from across each human Repli-Seq sample. To identify constitutive ERDs and LRDs (cERDs and cLRDs), ERDs and LRDs classified by Liu *et al* (Liu et al. 2015) across each cell type were evenly split into 1 kb intervals. If the classification of 12 out of 16 samples agreed across a certain 1 kb interval, we classified that region as belonging to a cERDs or cLRDs, depending the region’s majority classification of the 1 kb interval.

### DNase1 cluster identification and activity

DNase1 sites across 15 cell lines were found using DNase-seq and DNase-chip as part of the open chromatin synthesis dataset for ENCODE generated by Duke University’s Institute for Genome Sciences & Policy, University of North Carolina at Chapel Hill, University of Texas at Austin, European Bioinformatics Institute and University of Cambridge, Department of Oncology and CR-UK Cambridge Research Institute (Table S4) (ENCODE Project Consortium 2012). Regions where P-values of contiguous base pairs were below 0.05 were identified as significant DNase1 hypersensitive sites (ENCODE Project Consortium 2012). From this we extracted significant DNase1 hypersensitive sites from each sample and pooled them. DNase1 hypersensitive sites were then merged into DNase1 clusters. Cluster activity was calculated as the number of total overlapping pooled DNase1 hypersensitive sites. We also extracted intervals between adjacent DNase1 clusters to look for enrichment of retrotransposons at DNase1 cluster boundaries.

### Extraction of intergenic and intron intervals

hg19 RefSeq gene annotations obtained from UCSC genome browser were used to extract a set of introns and intergenic intervals (Table S5). RefSeq gene annotations were merged and intergenic regions were classified as regions between the start and end of merged gene models. We used the strandedness of gene model boundaries to classify adjacent intergenic region boundaries as upstream or downstream. We discarded intergenic intervals adjacent to gene models where gene boundaries were annotated as both + and − strand. Regions between adjacent RefSeq exons within a single gene model were classified as introns. Introns interrupted by exons in alternatively spliced transcripts and introns overlapped by other gene models were excluded. Upstream and downstream intron boundaries were then annotated depending on the strandedness of the gene they were extracted from.

### Interval boundary density of retrotransposons

Intervals were split in half and positions were reckoned relative to the feature adjacent boundary, where the feature was either a gene, exon, or DNase1 cluster (Fig. S2). To calculate the retrotransposon density at each position, we measured the fraction of bases at each position annotated as a retrotransposon. Next, we smoothed retrotransposon densities by calculating the mean and standard deviation of retrotransposon densities within an expanding window, where window size grows as a linear function of distance from the boundary. This made it possible to accurately compare the retrotransposon density at positions where retrotransposon insertions were sparse and density levels at each position fluctuated drastically. At positions with a high base pair density a small window was used and at positions with a low base pair density a large window was used. Expected retrotransposon density *p* was calculated as the total proportion of bases covered by retrotransposons across all intervals. Standard deviation at each position was calculated as 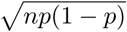, where *n* is the total number of bases at a given position.

### Interval size bias correction of retrotransposon densities

Interval boundary density is sensitive to retrotransposon insertion preferences into intervals of a certain size (Fig. S3). To determine interval size retrotransposon density bias, we grouped intervals according to size and measured the retrotransposon density of each interval size group. Retrotransposon density bias was calculated as the observed retrotransposon density of an interval size group divided by the expected retrotransposon density, where the expected retrotransposon density is the total retrotransposon density across all intervals. Next, using the intervals that contribute to the position depth at each position adjacent to feature boundaries, we calculated the mean interval size. From this we corrected retro-transposon density at each position by dividing the observed retrotransposon density by the retrotransposon density bias that corresponded with that position’s mean interval size.

### Software and data analysis

All statistical analyses were performed using R (R Core Team 2015) with the packages GenomicRanges (Lawrence et al. 2013) and rtracklayer (Lawrence et al. 2009). R scripts used to perform analyses can be found at: https://github.com/AdelaideBioinfo/retrotransposonAccumulation.

## Results

### Species selection and retrotransposon classification

We selected human, chimpanzee, rhesus macaque, mouse, rabbit, dog, horse and pig as representative placental species because of their similar non-LTR retrotransposon composition (Fig. S4-S5) and phylogenetic relationships. Retrotransposon coordinates were obtained from UCSC repeat masker tables and the online repeat masker database (Rosenbloom et al. 2015; Smit et al. 1996). We grouped non-LTR retrotransposon families according to repeat type and period of activity as determined by genome-wide defragmentation (Giordano et al. 2007). Retrotransposons were placed into the following groups; new L1s, old L1s, new SINEs and ancient elements (for families in each group see Fig. S5). New L1s and new SINEs are retrotransposon families with high lineage specificity and activity, while old L1s and ancient elements (SINE MIRs and LINE L2s) are retrotransposon families shared across taxa. We measured sequence similarity within retrotransposon families as percentage mismatch from family consensus sequences (Bao et al. 2015). We found that more recent lineage-specific retrotransposon families had accumulated a lower percentage of substitutions per element than older families (Fig. S6-S13). This confirmed that our classification of retrotransposon groups agreed with ancestral and lineage-specific periods of retrotransposon activity.

### Genomic distributions of retrotransposons

To analyse the large scale distribution of retrotransposons, we segmented each species genome into adjacent 1 Mb regions, tallied retrotransposon distributions, performed principal component analysis (PCA) and pairwise correlation analysis (see methods). For PCA, our results showed that retrotransposon families from the same group tended to accumulate in the same genomic regions. We found that each individual retrotransposon group was usually highly weighted in one of the two major principal components (PC1 and PC2) (Fig. 2). Depending on associations between PCs and particular retrotransposon families we identified PC1 and PC2 as either the “lineage-specific PC” or the “ancestral PC”. Along the lineage-specific PC, new SINEs and new L1s were highly weighted, where in all species new SINEs were enriched in regions with few new L1s. Alternatively, along the ancestral PC, old L1s and ancient elements were highly weighted, where in all species except mouse — where ancient elements and old L1s were co-located — ancient elements were enriched in regions with few old L1s (Fig. 2–3a, S14). The discordance observed in mouse probably resulted from the increased genome turnover and rearrangement seen in the rodent lineage potentially disrupting the distribution of ancestral retrotransposon families (Murphy et al. 2005; Capilla et al. 2016). In addition, the genome-wide density of ancestral retrotransposons in mouse was particularly low compared to our other species (Fig. S4-S5). However, as the relationship between mouse lineage-specific new retrotransposons is maintained, this discordance does not impact on downstream analyses. These results show that most genomic context associations between retrotransposon families are conserved across our sample species.

**Figure 2.**
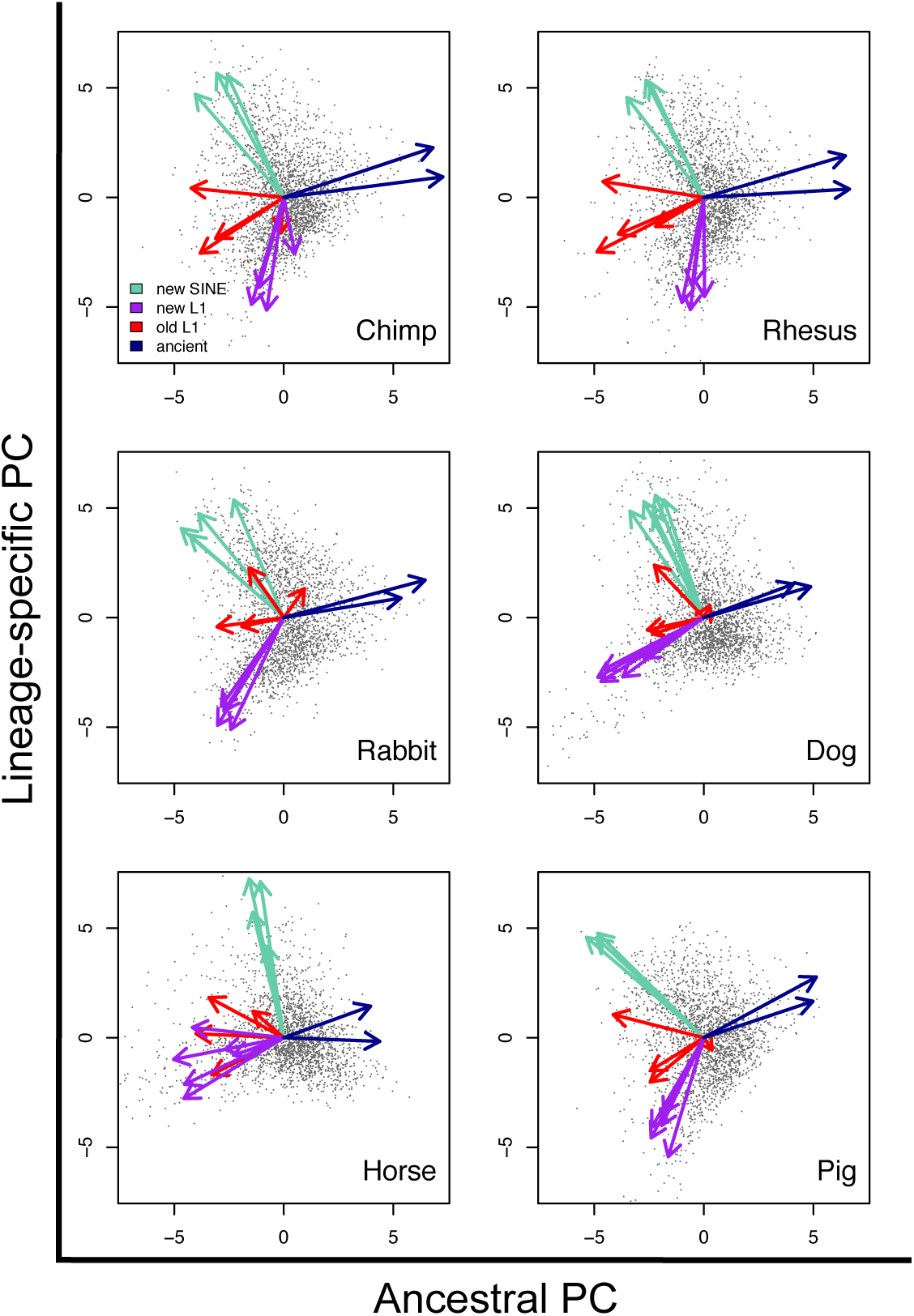
Similar genomic distributions of retrotransposons across mammals. Principal Component 1 and Principal Component 2 of non-human and non-mouse genome retrotransposon content, each vector loading has been coloured according to the retrotransposon group it represents. Principal components have been renamed according to the retrotransposon group whose variance they principally account for.

### Retrotransposon accumulation and chromatin environment

In human and mouse, LINEs and SINEs differentially associate with distinct chromatin environments (Ashida et al. 2012). To determine how our retrotransposon groups associate with chromatin accessibility, we obtained ENCODE generated human cell line Repli-Seq data and mouse cell line Repli-ChIP data from the UCSC genome browser (ENCODE Project Consortium 2012; Yue et al. 2014). Repli-Seq and Repli-CHiP both measure the timing of genome replication during S-phase, where accessible euchromatic domains replicate early and inaccessible heterochromatic domains replicate late. Across our segmented genomes, we found a high degree of covariation between genome-wide mean replication timing and lineage-specific PC scores (Fig. 3a), new SINEs associated with early replication and new L1s associated with late replication. In addition, by splitting L1s into old and new groups, we showed a strong association between replication timing and retrotransposon age that was not reported in previous analyses (Pope et al. 2014). These results are probably not specific to a particular cell line, since genome-wide replication timing patterns are mostly highly correlated across cell lines from either species (Table S6). Moreover, early and late replicating domains from various human cell lines exhibit a high degree of overlap (Fig. S15). To confirm that lineage-specific retrotransposon accumulation associates with replication timing, we analysed retrotransposon accumulation at the boundaries of previously identified replication domains (RDs) (Liu et al. 2015). We focused primarily on early replicating domain (ERD) boundaries rather than late replicating domain (LRD) boundaries because ERD boundaries mark the transition from open chromatin states to closed chromatin states and overlap with topologically associated domain (TAD) boundaries (Pope et al. 2014). Consistent with our earlier results, significant density fluctuations at ERD boundaries were only observed for new L1s and new SINEs (Fig. 3b). Because RD timing and genomic distributions of clade-specific retrotransposons are both largely conserved across human and mouse (Ryba et al. 2010; Yaffe et al. 2010), these results suggest that the relationship between retrotransposon accumulation and RD timing may be conserved across mammals.

**Figure 3.**
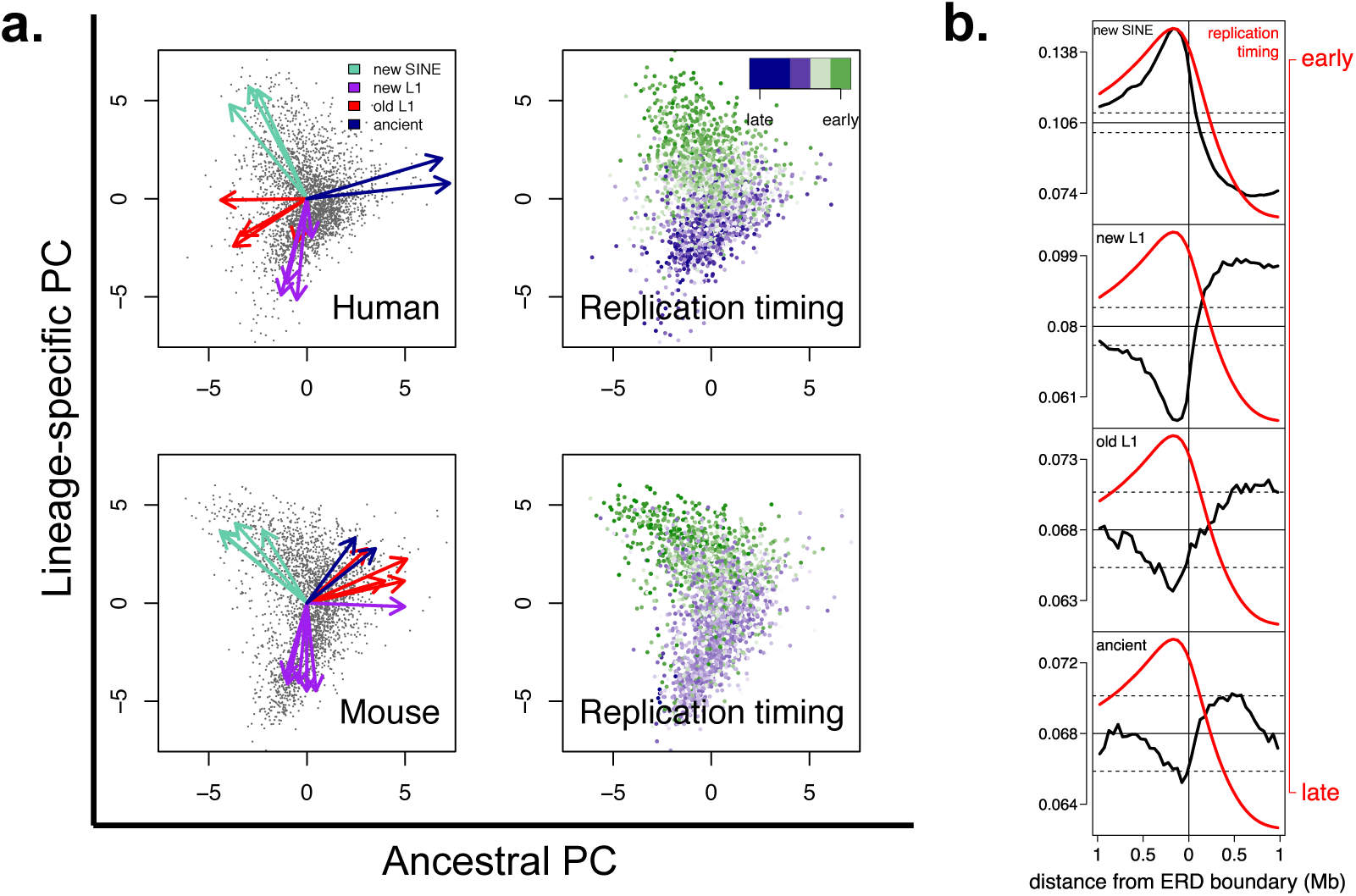
**a**, PCA of human and mouse retrotransposon content and mean genome replication timing in human HUVEC cells and mouse EpiSC-5 cells. **b**, Retrotransposon density per non-overlapping 50 kb intervals from a pooled set of ERD boundaries across all 16 human cell lines. Black dashed lines indicate 2 standard deviations from the mean (solid horizontal black line). Red line indicates mean replication timing across all samples.

### The genomic distribution of retrotransposons is conserved across species

Our earlier results showed that the genomic distribution of retrotransposons is similar across species (Fig. 2). To determine whether our observations resulted from retrotransposon insertion into orthologous regions, we humanised segmented genomes of non-human species. Humanisation, began with a segmented human genome, a segmented non-human mammalian genome, and a set of pairwise alignments between both species. Using the pairwise alignments we calculated the percentage of nucleotides from each human segment that aligned to a specific non-human segment and vice-versa. This made it possible to remodel the retrotransposon content of each non-human genome segment within the human genome and essentially humanise non-human mammalian genomes (Fig. 1) (see methods). To test the precision of our humanisation process, we used the Kolmogorov-Smirnov test to compare the humanised retrotransposon density distribution of a specific retrotransposon family, to the non-humanised retrotransposon density distribution of that same retrotransposon family (Fig. S1). If the Kolmogorov-Smirnov test returned a low P-value, this suggested that the humanisation process for a given retrotransposon family had a low level of precision. Therefore, to increase our precision we used a minimum mapping fraction threshold to discard genomic segments that had only had a small amount of aligning regions between each genome. The motivation behind this was that genomic segments with a small amount of aligning sequence were the ones most likely to inaccurately represent non-human retrotransposon genomic distributions when humanised. However, it is important to note that our increase in precision requires a trade-off in accuracy. By discarding genomic segments below a certain threshold we sometimes removed a significant fraction of our non-human genomes from the analysis. In addition, this approach disproportionately affected retrotransposons such as new L1s, as they were most enriched in segments with a small amount of aligning regions between each genome (Fig.S16-S17). To overcome this, we humanised each non-human genome at minimum mapping fraction thresholds of 0, 10, 20, 30, 40 and 50 percent and recorded the percentage of the genome that remained. We found that most retrotransposon families were precisely humanised at a minimum mapping fraction threshold of 10%. In non-human species where humanisation was most precise, a minimum mapping fraction threshold of 10% resulted in greater than 90% of the human and non-human genome remaining in the analysis (Fig. 4, S18-S24). After humanising each non-human genome, we performed pairwise correlation analysis (see methods) between the genomic distributions of each humanised and human retrotransposon family. Our results showed that retrotransposon families in different species that were identified as the same group showed relatively strong correlations, suggesting that they accumulated in regions with shared common ancestry (Fig. 4, S18-S24). Next, we assessed the level of conservation of retrotransposon accumulation patterns across all of our species. For each retrotransposon group in each humanised genome, we identified the top 10% retrotransposon dense genome segments. We found that when these segments were compared with the human genome, there was a relativity high degree of overlap (Fig. 5a-b). These results suggest that lineage-specific retrotransposon accumulation may follow an ancient conserved mammalian genome architecture.

**Figure 4.**
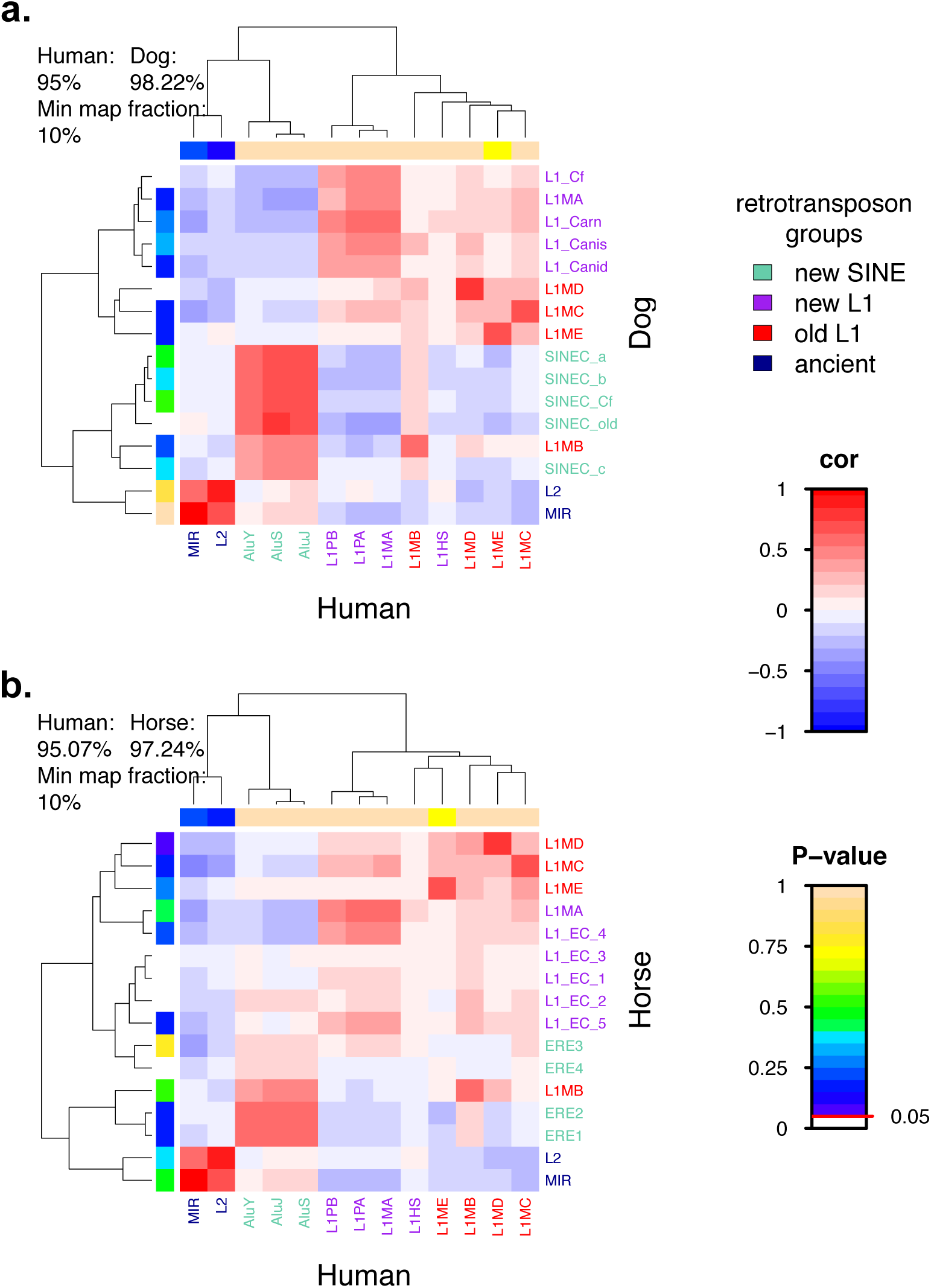
Genome-wide spatial correlations of humanised retrotransposon families. Heatmap colours represent Pearson’s correlation coefficient for genomic distributions between humanised **a**, dog and human retrotransposon families, and humanised **b**, horse and human retrotransposon families. Values at the top left of each heatmap reflect the proportion of each genome analysed after filtering at a 10% minimum mapping fraction threshold (Fig. 1a). Dog and horse P-values represent the effect of humanising on filtered non-human retrotransposon density distributions (Fig. 1e). Human P-values represent the effect of filtering on the human retrotransposon density distributions (Fig. 1f).

**Figure 5.**
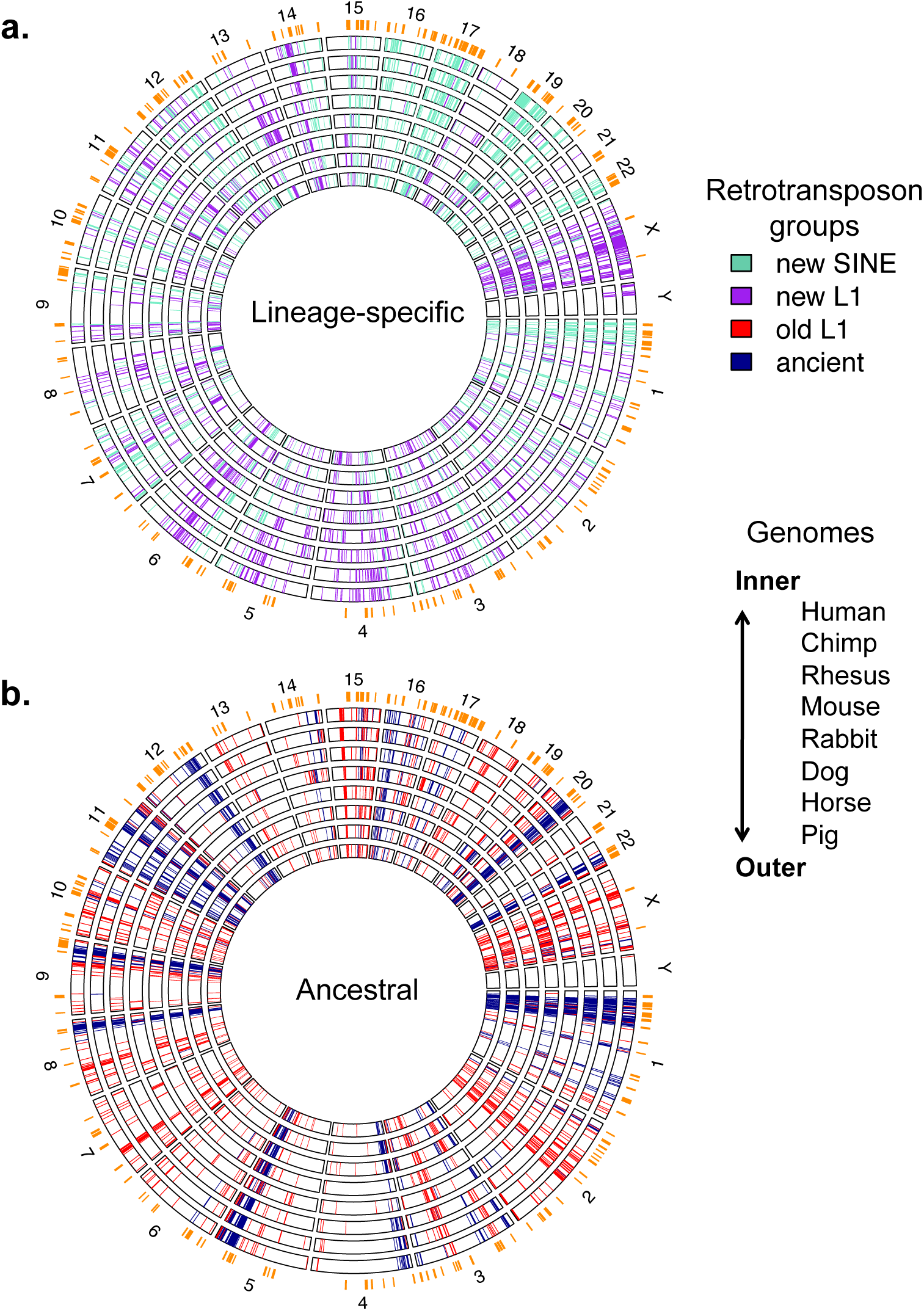
Retrotransposon accumulation patterns are conserved across mammals. **a**, Top 10% of genome segments based on retrotransposon density of new SINEs and new L1s. **b**, Top 10% of genome segments based on retrotransposon density of ancient elements and old L1s. In both **a** and **b**, segments for non-human genomes were ranked according to their humanised values. Large ERDs (> 2 Mb) from HUVEC cells are marked in orange.

### Retrotransposon insertion in open chromatin surrounding regulatory elements

Retrotransposons preferentially insert into open chromatin, yet open chromatin usually overlaps gene regulatory elements. As stated above, this creates a fundamental evolutionary conflict for retrotransposons; their immediate replication may be detrimental to the overall fitness of the genome in which they reside. To investigate retrotransposon insertion/accumulation dynamics at open chromatin regions, we analysed DNase1 hypersensitive activity across 15 cell lines in both ERDs and LRDs. DNase1 hypersensitive sites obtained from the UCSC genome browser (ENCODE Project Consortium 2012) were merged into DNase1 clusters and DNase1 clusters overlapping exons were excluded. As replication is sometimes cell type-specific we also constructed a set of constitutive ERDs and LRDs (cERDs and cLRDs) (see methods). Based on previous analyses, cERDs and cLRDs likely capture RD states present during developmental periods of heritable retrotransposition (Rivera-Mulia et al. 2015). Our cERDs and cLRDs capture approximately 50% of the genome and contain regions representative of genome-wide intron and intergenic genome structure (Fig. S25). In both cERDs and cLRDs, we measured DNase1 cluster activity by counting the number of DNase1 peaks that overlapped each cluster. We found that DNase1 clusters in cERDs were much more active than DNase1 clusters in cLRDs (Fig. 6a). Next, we analysed retro-transposon accumulation both within and at the boundaries of DNase1 clusters. Consistent with disruption of gene regulation by retrotransposon insertion, non-ancient retrotransposon groups were depleted from DNase1 clusters (Fig. 6b). Intriguingly, ancient element density in DNase1 clusters remained relatively high, suggesting that some ancient elements may have been exapted. At DNase1 cluster boundaries after removing interval size bias (Fig. S26-S27) (see methods), retrotransposon density remained highly enriched in cERDs and close to expected levels in cLRDs (Fig. 6c). This suggests that chromatin is likely to be open at highly active cluster boundaries where insertion of retrotransposons is less likely to disrupt regulatory elements. These results are consistent with an interaction between retrotransposon insertion, open chromatin and regulatory activity, where insertions into open chromatin only persist if they do not interrupt regulatory elements.

**Figure 6.**
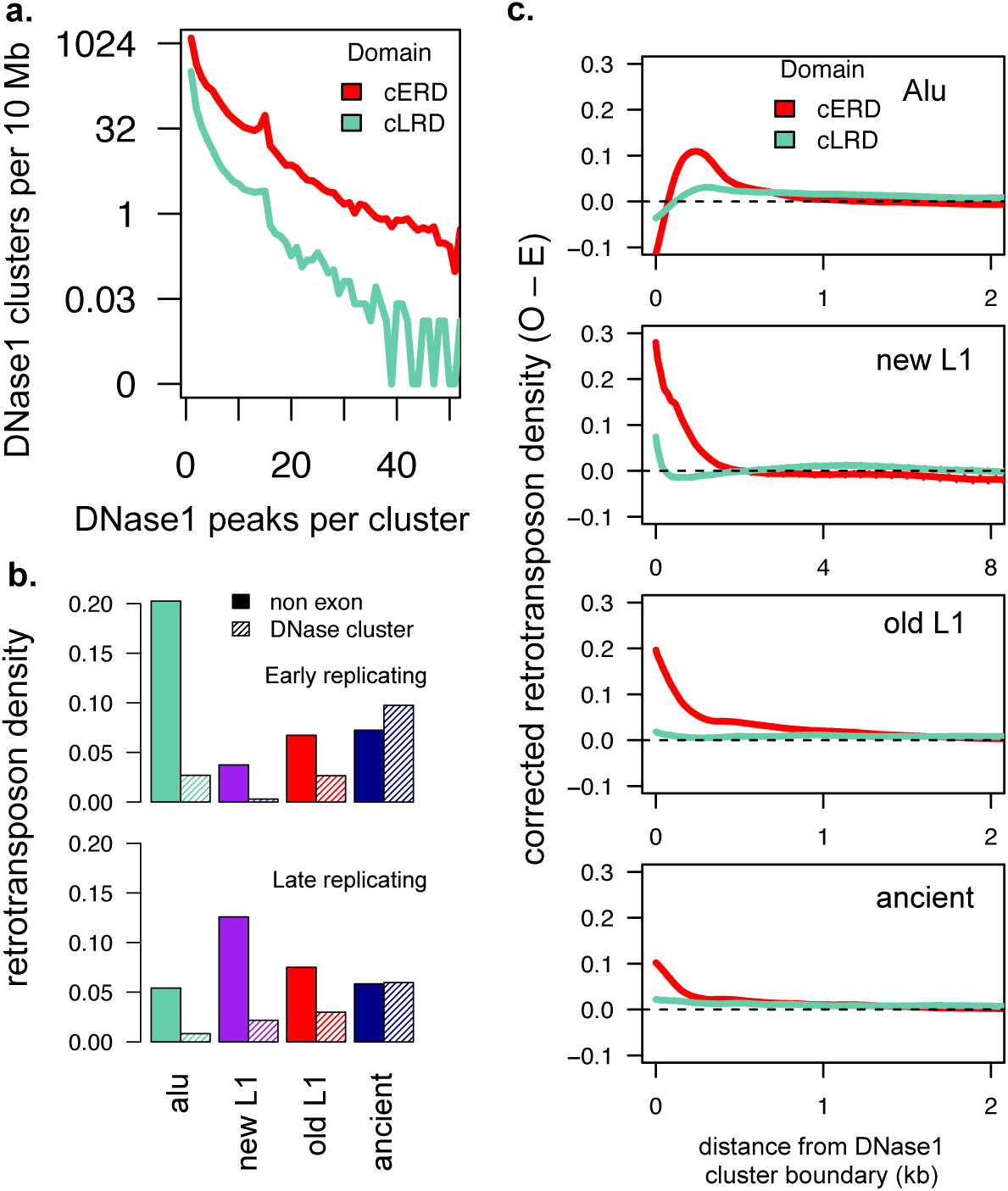
Retrotransposon accumulation occurs in open chromatin near regulatory regions. **a**, The activity of DNase1 clusters in cERDs and cLRDs. DNase1 clusters were identified by merging DNase1 hypersensitive sites across 15 tissues. Their activity levels were measured by the number of DNase1 hypersensitive sites overlapping each DNase1 cluster. **b**, Retrotransposon density of non-exonic regions and DNase1 clusters in cERDs and cLRDs. **c**, Observed minus expected retrotransposon density at the boundary of DNase1 clusters corrected for interval size bias (see methods). Expected retrotransposon density was calculated as each group’s non-exonic total retrotransposon density across cERDs and cLRDs. A confidence interval of 3 standard deviations from expected retrotransposon density was also calculated, however the level of variation was negligible.

### Retrotransposon insertion size and regulatory element density

L1s and their associated SINEs differ in size by an order of magnitude, retrotranspose via the L1-encoded chromatin-sensitive L1ORF2P and accumulate in compositionally distinct genomic domains (Cost et al. 2001; Baillie et al. 2011). This suggests that retrotransposon insertion size determines observed accumulation patterns. L1 and *Alu* insertions occur via target-primed reverse transcription which is initiated at the 3′ end of each element. With L1 insertion, this process often results in 5′ truncation, causing extensive insertion size variation and an over representation of new L1 3′ ends, not seen with *Alu* elements (Fig. 7a). When we compared insertion size variation across cERDs and cLRDs we observed that smaller new L1s were enriched in cERDs and *Alu* elements showed no RD insertion size preference (Fig. 7b). The effect of insertion size on retrotransposon accumulation was estimated by comparing insertion rates of each retrotransposon group at DNase1 cluster boundaries in cERDs and cLRDs. We found that *Alu* insertion rates at DNase1 cluster boundaries were similarly above expected levels both in cERDs and cLRDs (Fig. 7c), whereas new L1 insertion rates at DNase1 cluster boundaries were further above expected levels in cERDs than cLRDs (Fig. 7d). By comparing the insertion rate of new L1s — retrotransposons that exhibited RD specific insertion size variation — we observed a negative correlation between element insertion size and gene/regulatory element density. Thus smaller elements, such as *Alu* elements, accumulate more in cERDs than do larger elements, such as new L1s, suggesting that smaller elements are more tolerated.

**Figure 7.**
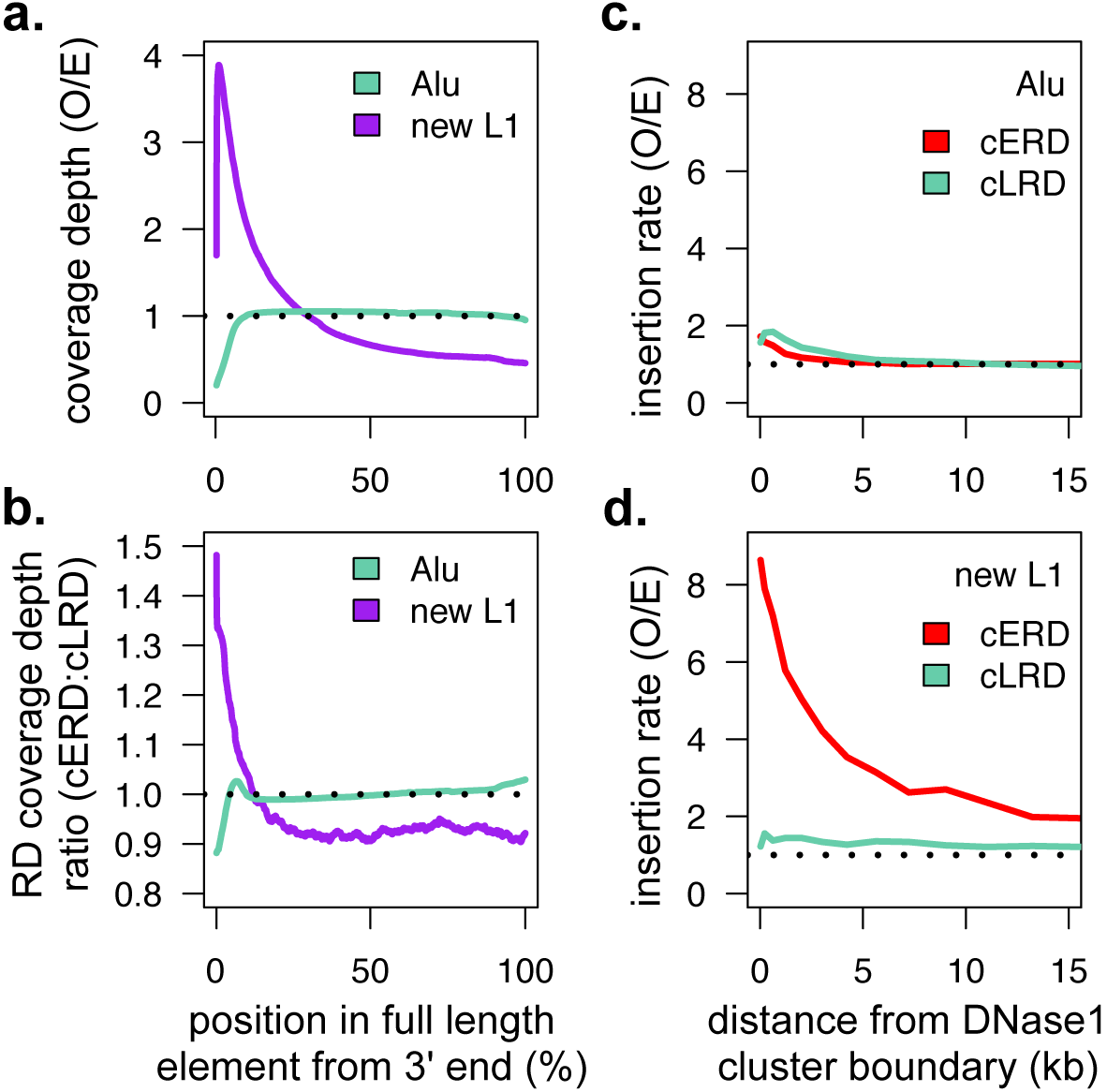
Retrotransposon insertion size is inversely proportional to local regulatory element density. **a**, Observed to expected ratio of retrotransposon position coverage depth measured from consensus 3′ end. Expected retrotransposon position coverage depth was calculated as total retrotransposon coverage over consensus element length. We used 6 kb as the consensus new L1 length and 300 bp as the consensus *Alu* length. **b**, New L1 and *Alu* position density ratio (cERDs:cLRDs). **c**, *Alu* and **d**, new L1 observed over expected retrotransposon insertion rates at DNase1 cluster boundaries in cERDs and cLRDs. Insertion rates were measured by prevalence of 3′ ends and expected levels were calculated as the per Mb insertion rate across cERDs and cLRDs.

### Retrotransposon insertion within gene and exon structures

Regulatory element organisation is largely shaped by gene and exon/intron structure which likely impacts the retrotransposon component of genome architecture. Therefore, we analysed retrotransposons and DNase1 clusters (exon-overlapping and exon non-overlapping) at the boundaries of genes and exons. Human RefSeq gene models were obtained from the UCSC genome browser and both intergenic and intronic regions were extracted (Table S5). At gene (Fig. 8a) and exon (Fig. 8b) boundaries, we found a high density of exon overlapping DNase1 clusters and depletion of retrotransposons. This created a depleted retrotransposon boundary zone (DRBZ) specific for each retrotransposon group, a region extending from the gene or exon boundary to the point where retrotransposon levels begin to increase. The size of each DRBZ correlated with the average insertion size of each retrotransposon group, consistent with larger retrotransposons having a greater capacity to disrupt important structural and regulatory genomic features. We also found that in cERDs the 5′ gene boundary *Alu* DRBZ was larger than the 3′ gene boundary *Alu* DRBZ. This difference was associated with increased exon overlapping DNase1 cluster density at 5′ gene boundaries in cERDs (Fig. 8a), emphasising the importance of evolutionary constraints on promoter architecture. For ancient elements, their retrotransposon density at approximately 1 kb from the 5′ gene boundary, when corrected for interval size bias, was significantly higher than expected. This increase is consistent with exaptation of ancient elements into regulatory roles (Lowe et al. 2007) (Fig. S28-S31). Moreover, the density peak corresponding to uncorrected ancient elements also overlapped with that of exon non-overlapping DNase1 clusters (Fig. 8a). Collectively, these results demonstrate the evolutionary importance of maintaining gene structure and regulation and how this in turn has canalised similar patterns of accumulation and distribution of retrotransposon families in different species over time.

**Figure 8.**
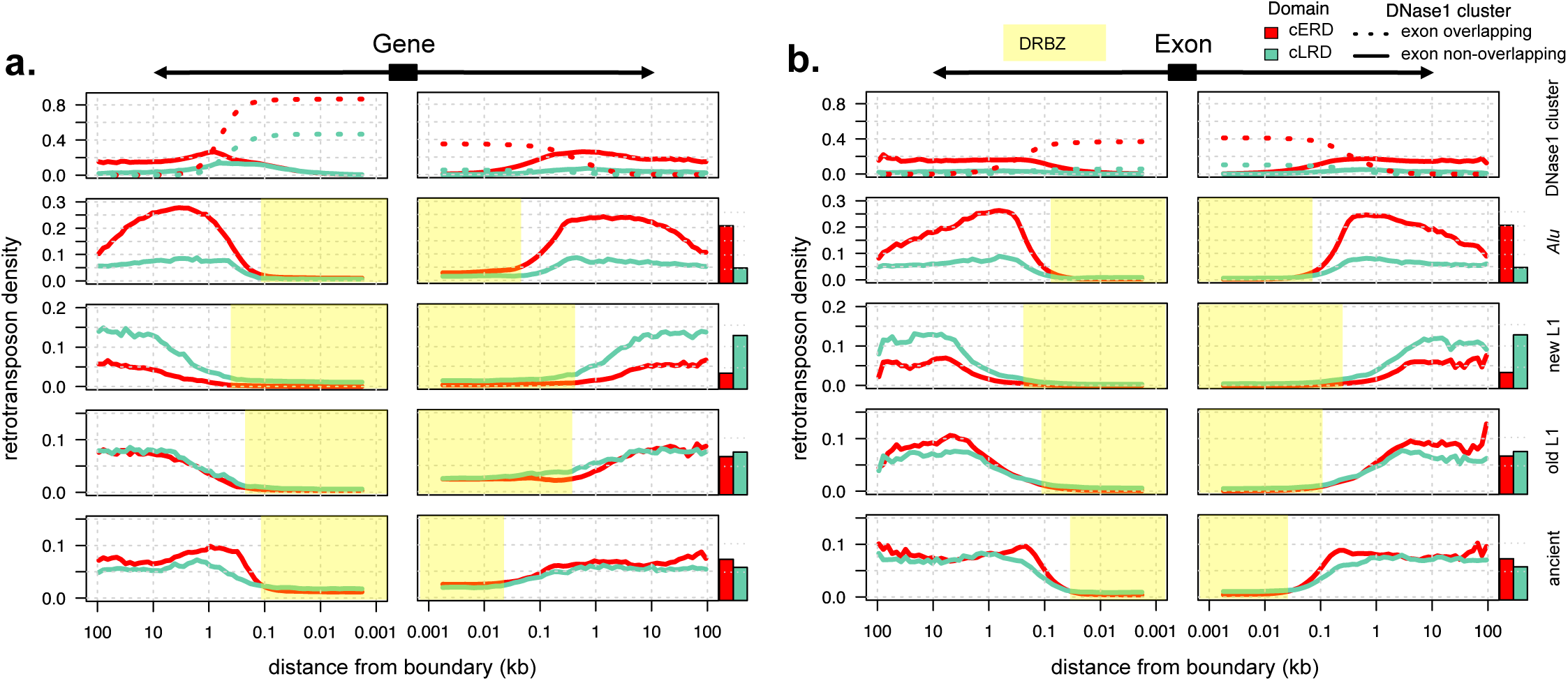
Retrotransposon accumulation within intergenic and intronic regions correlates with the distribution of DNase1 clusters. Density of DNase1 clusters, and retrotransposons at each position upstream and downstream of genes and exons in **a**, intergenic and **b**, intronic regions. For DNase1 clusters, dotted lines represent exon overlapping clusters and solid lines represent clusters that do not overlap exons. For retrotransposons, solid lines represent the uncorrected retrotransposon density at exon and gene boundaries. Bar plots show expected retrotransposon density across cERDs and cLRDs. Highlighted regions outline DRBZs, regions extending from the gene or exon boundary to the point where retrotransposon levels begin to increase.

## Discussion

### A conserved architectural framework shapes the genomic distribution of ancestral retrotransposons

The majority of divergence between our sample species has taken place over the last 100 million years. Throughout this time period many genomic rearrangements have occurred, causing a great deal of karyotypic variation. However, we found that the genomic distributions of ancestral elements remained conserved. The evolutionary forces preserving the ancestral genomic distributions of these elements remain unclear.

One suggestion is that ancestral elements play essential roles in mammalian organisms. Our results in Fig. 6b and 8a suggest that ancient elements have been exapted. Their accumulation within open chromatin sites is consistent with their roles as *cis*-regulatory element, such as MIR elements that perform as TFBSs and enhancers (Bourque et al. 2008; Jjingo et al. 2014). Similarly, L1s also carry binding motifs for DNA-binding proteins. L1 elements that were active prior to the boreoeutherian ancestor bind a wide variety of KRAB zinc-finger proteins (KZFPs), most of which have unknown functions (Imbeault et al. 2017). In terms of genome structural roles, some human MIR elements have been identified as insulators, separating open chromatin regions from closed chromatin regions (Wang et al. 2015). While these MIR insulators function independently of CTCF binding, their mechanism of action remains largely unknown. Despite this, when a human MIR insulator was inserted into the zebrafish genome it was able to maintain function (Wang et al. 2015). This suggests that MIR insulators recruit a highly conserved insulator complex and maintain insulator function across the mammalian lineage. Collectively, these findings identified a number of examples where ancestral elements are associated with important biological roles. This may suggest that genomic distributions of ancestral elements are conserved across mammals because they play conserved biological roles across mammals. However, it is necessary to draw a distinction between evolutionary conservation of an ancient functional element and evolutionary conservation of large-scale genomic distributions of retrotransposons. This is important because for most of our sample species, ancient elements and old L1s each occupy approximately 7% of each of their genomes (Fig. S4). Compared to the 0.04% of the human genome that is comprised of transposable elements under purifying selection (Lowe et al. 2007), this suggests that the vast majority of ancestral elements may not actually play conserved roles in mammalian biology.

Rather than ancestral elements playing a conserved role in genome maintenance, their genomic distributions may instead remain conserved as a consequence of evolutionary dynamics occurring at higher order levels of genome architecture. TADs have been identified as a fundamental unit of genome structure, they are approximately 900 kb in length and contain highly self interacting regions of chromatin (Dixon et al. 2012). Despite large-scale genomic rearrangements, the boundaries between TADs have remained conserved across mammals (Dixon et al. 2012). An analysis involving rhesus macaque, dog, mouse and rabbit, identified TAD boundaries at the edge of conserved syntenic regions associating with evolutionary breakpoints between genomic rearrangements (Rudan et al. 2015). This suggests that genome rearrangements occur primarily along TAD boundaries leaving TADs themselves largely intact. Similarly, TAD architecture could also be the driving force behind the observed frequent reuse of evolutionary breakpoints throughout mammalian genome evolution (Murphy et al. 2005). Together these findings suggest that TADs form part of a conserved evolutionary framework whose boundaries are sensitive to genomic rearrangements. Therefore, the current observed genomic distributions of ancestral retrotransposons reflects mostly ancestral retrotransposons that inserted within TADs rather than at their boundaries. This is because elements that accumulated near TAD boundaries were most likely lost through recurrent genomic rearrangements and genome turnover.

Another example supporting the idea that conserved genomic distributions are shaped by a conserved architectural evolutionary framework can be found in the rodent lineage. Rodents have experienced rates of genome reshuffling two orders of magnitude greater than other mammalian lineages (Capilla et al. 2016). This has caused rodent genomes to contain a higher number of evolutionary breakpoints, many of which are rodent-specific (Capilla et al. 2016). From our analysis we found that old L1s and ancient elements each occupied only 1% of the mouse genome (Fig. S4), with similar levels of ancient elements within the rat genome (Gibbs et al. 2004). Compared to our other species where the genomes are approximately 7% ancient elements and old L1s each (S4), rodent genomes are significantly depleted of ancestral elements. Together, these findings show a negative correlation between ancestral retrotransposon content and rate of genome rearrangements, suggesting that increased rates of genome rearrangements can strongly impact the genomic distributions of ancestral retrotransposons. In addition, the large number of rodent specific evolutionary breakpoints may explain why the genomic distribution of ancestral elements in mouse is discordant with our other species. Specifically, ancient elements and old L1s in mouse accumulated in similar regions, whereas in each of our other species ancient elements and old L1s accumulated in almost opposite regions as defined by PC1 (Fig. 2, 3a).

### Conserved genome architecture drives the accumulation patterns of lineage-retrotransposons

Across mammals, lineage-specific retrotransposons are responsible for the vast majority of lineage-specific DNA gain (Kapusta et al. 2017). Throughout our sample-species we found that new SINEs and new L1s independently accumulated in similar regions in different species. These results suggest there is a high degree of conservation surrounding their insertion mechanisms and genomic environments. Since, L1 conservation in mammals is well documented in the literature and our new SINE families all replicate using L1 machinery, mainly we spend this section discussing the role of conserved genome architecture (Ivancevic et al. 2016; Vassetzky and Kramerov 2013).

Earlier, we discussed the importance of TADs and how they form a fundamental component of conserved genome architecture. This same architectural framework may also shape the accumulation pattern of lineage specific retrotransposons. TAD boundaries separate the genome into regions comprised of genes that are largely regulated by a restricted set of nearby enhancers. Moreover, TADs are subject to large-scale changes in chromatin structure, where individual TADs are known to switch between open and closed chromatin states in a cell type-specific manner (Dixon et al. 2012). One method of capturing shifts in chromatin state between TADs is to measure genome-wide replication timing (Pope et al. 2014). This is because replication timing associates with the genomes accessibility to replication machinery. Accessible regions that comprise an open chromatin structure replicate early while inaccessible regions with a closed chromatin structure replicate late. Genome-wide replication timing follows a domain-like organisation, where large contiguous regions either replicate at earlier or later stages of mitosis. Importantly, ERD boundaries directly overlap TAD boundaries, supporting the notion that TADs are also fundamental units of large-scale chromatin state organisation (Pope et al. 2014). Previously, LINE and SINE accumulation patterns were associated with TAD and RD genome architecture, where LINEs were enriched in LRDs and SINEs were enriched in ERDs (Hansen et al. 2010; Rivera-Mulia et al. 2015; Pope et al. 2014; Ashida et al. 2012). Unlike our analysis, these earlier studies decided not to separate LINEs into ancestral and lineage-specific families. Despite this difference, Fig. 3 shows that our results are consistent with earlier analyses, except for our observation that only lineage-specific retrotransposon families are associated with replication timing. Therefore, by separating L1s and SINEs according to period of activity, we observed much stronger associations between replication timing and retrotransposon accumulation than previously reported (Pope et al. 2014; Ashida et al. 2012). Since replication timing and boundaries between TADs and RDs are conserved across mammalian species (Ryba et al. 2010; Yaffe et al. 2010; Pope et al. 2014; Dixon et al. 2012), our results suggest that domain-level genome architecture likely plays a role in shaping conserved lineage-specific retrotransposon accumulation patterns.

While our species genomes are conserved at a structural level, conserved patterns of lineage-specific retrotransposon accumulation can have significant evolutionary impacts. new SINEs accumulate in ERDs which tend to be highly active gene-rich genomic regions. However, despite the fact that all of our new SINE families follow L1 mediated replication, they stem from unique origins. For example, Primate-specific *Alu* elements are derived from 7SL RNA and carnivora-specifc SINEC elements are dervided form tRNA (Quentin 1994; Coltman and Wright 1994). Due to their large-scale accumulation patterns this means that new SINEs in mammalian genomes simultaneously drive convergence in genome architecture and divergence in genome sequence composition. This is especially important because SINEs are also a large source of evolutionary innovation for gene regulation. In human, various individual *Alu* elements have been identified as bona fide enhancers with many more believed to be proto-enhacers serving as a repertoire for birth of new enhancers (Su et al. 2014). Similarly, in dog, mouse and opossum, lineage specific SINEs carry CTCF binding sites and have driven the expansion of species-specific CTCF binding patterns (Schmidt et al. 2012).

Like new SINEs, new L1s also accumulate in similar regions in different species. However, unlike new SINEs, lineage-specific mammalian L1 elements most likely stem from a common ancestor (Furano et al. 2004). This means that individual new L1 elements in different species are more likely than species-specific SINEs to share similar sequence composition (Ivancevic et al. 2016). Therefore, LRDs, which are enriched for new L1s, may show higher levels of similarity for genome sequence composition than ERDs, which are enriched for new SINEs. Considering results from genome-wide alignments between mammals, this may be counter intuitive, mainly because the surrounding sequence in new L1 enriched regions exhibits poor sequence conservation (Fig. S16-S17). However, it is important to realise that similar sequence composition is not the same as sequence conservation itself, especially at the level of mammalian genome architecture. Sequence composition refers to the kinds of sequences in a particular region rather than the entire sequence of the region itself. For example, binding sites for the same transcription factor in different species are sometimes located in similar regions yet differ in position relative to their target genes (Kunarso et al. 2010). So while genome-wide alignments may suggest low levels of genome conservation or high levels of turnover, sequence composition within these regions remains similar and can still be indicative of conserved function. Therefore with the accumulation of new L1s after species divergence, it is likely that sequence conservation decreases at a much faster rate than compositional similarity. For new L1s enriched in similar regions in different species, this may have important functional consequences. Recently, highly conserved ancient KZFPs were discovered to bind to members of both old and new L1 families in human (Imbeault et al. 2017). This suggests that new L1s in humans may be interchangeable with old L1s and play important roles in highly conserved gene regulatory networks. Therefore, because new L1s in different species share similar sequences and their accumulation patterns are also conserved, new L1s may actively preserve ancient gene regulatory networks across the mammalian lineage.

### A chromatin based model of retrotransposon accumulation

Analysis of repetitive elements in mammalian genome sequencing projects has consistently revealed that L1s accumulate in GC-poor regions and their mobilised SINEs accumulate in GC-rich regions (Lander et al. 2001; Gibbs et al. 2004; Chinwalla et al. 2002). Our results were consistent with this and showed that accumulation patterns of new SINEs and new L1s were conserved across species and corresponded with distinct genomic environments. Since these elements both replicate via the same machinery, their accumulation patterns are most likely shaped by how insertion of each element type interacts with its immediate genomic environment. The current model of retrotransposon accumulation begins with random insertion, constrained by local sequence composition, followed by immediate selection against harmful insertions (Graham and Boissinot 2006; Gasior et al. 2007; Kvikstad and Makova 2010). During early embryogenesis or in the germline, it is believed retrotransposons in individual cells randomly insert into genomic loci that contain a suitable insertion motif. Because this process is assumed to be random, new insertions can occasionally interrupt essential genes or gene regulatory structures. These insertions are usually harmful, causing the individual cell carrying them to be quickly removed from the population. This process of purifying selection prevents harmful insertions from being passed down to the next generation and plays a large role in shaping retrotransposon accumulation patterns. According to this model, because of their size difference L1s are considered to have a more harmful impact on nearby genes and gene regulatory structures than SINEs. New L1 insertion into GC-rich regions, which are also gene-rich, are more likely to cause harm than if new SINEs inserted into those same regions. Therefore, new L1s are evolutionary purged from GC rich regions causing them to become enriched in gene-poor AT-rich regions. While this model is simple, it fails to take into account the impact of chromatin structure that constrains retrotransposon insertion preference. Therefore, we decided to analyse retrotransposon accumulation at the level of large-scale chromosomal domains and fine-scale open chromatin sites.

Our results showed that lineage-specifc retrotransposons accumulated at the boundaries of open chromatin sites. This was particularly striking as it appeared to reconcile insertion into open chromatin with the risk of disrupting regulatory elements. Single cell analysis has shown somatic retrotransposition events correlate with preferable insertion into open chromatin sites or within actively expressed genes (Klawitter et al. 2016; Upton et al. 2015; Baillie et al. 2011). However, because open chromatin usually surrounds regulatory elements these kinds of insertions can be a major cause of genetic disease (Wimmer et al. 2011). Therefore, retrotransposons accumulate in open chromatin regions where their insertion is less likely to disrupt regulatory elements. We further demonstrated the impact of retrotransposon insertion by considering element insertion size. Our results showed that shorter L1s were much more likely to insert close to open chromatin sites surrounding regulatory elements than larger L1s. This suggested that L1 insertions were much more likely than *Alu* insertions to impact on gene regulatory structures due to their larger insertion size. At this point, it should be noted that chromatin state can be highly dynamic, switching between open and closed states depending on cell type (ENCODE Project Consortium 2012). Importantly, heritable retrotransposon insertions typically occur during embryogenesis or within the germline. However, chromatin state data for these developmental stages and tissue samples was unavailable. To overcome this limitation we aggregated data from a range of biological contexts. The underlying assumption behind this strategy was that open chromatin sites found in at least one cell likely contain regulatory elements that may be reused in another cell type. Theretofore, by using this strategy, we increased the probability of capturing chromosomal domain structures and regulatory element sites present in embryonic and germline cell states.

An important aspect of both our refined model and the current model of retrotransposon accumulation is the immediate evolutionary impact of retrotransposon insertions. Specifically, at what rate do embryonic and germline retrotransposition events occur and what proportion of these events escape purifying selection? Answering this question is a challenging task primarily limited by the availability of samples at the correct developmental time periods. Ideally we would require genome sequencing data from a large population of germline or embryonic cells derived from a similar genetic background. Given that data, we could identify new insertions before they have undergone selection and compare their retrotransposition rates to retrotransposition rates inferred from population data. Alternatively, retrotransposition rates have been measured in somatic cells and stem-cell lines. In hippocampal neurons and glia, L1 retrotransposition occurs at rates of 13.7 and 6.5 events per cell, where in human induced pluripotent stem cells retrotransposition rates are approximately 1 event per cell (Klawitter et al. 2016; Upton et al. 2015). In neurons, L1s insertions were enriched in neuronally expressed genes and in human induced pluripotent stem cells, L1s were found to insert near transcription start sites, disrupting the expression of some genes (Klawitter et al. 2016; Upton et al. 2015; Baillie et al. 2011). This suggests L1s are particularly active in humans, able to induce a large amount of variation and disrupt gene regulation and function. It is also important to note that the estimated L1 heritable retrotransposition rate is approximately one event per 95 to 270 births (Ewing and Kazazian 2010), suggesting that many insertions are removed from the germline cell population. For *Alu* elements this rate is much greater, *Alu* elements are estimated to undergo heritable retrotransposition at a rate of one event per 20 births (Cordaux et al. 2006). These findings support the notion that the majority of retrotransposon insertions are likely to be evolutionarily purged from the genome.

In summary, by analysing open chromatin sites, we found that 1) following preferential insertion into open chromatin domains, retrotransposons were tolerated adjacent to regulatory elements where they were less likely to cause harm; 2) element insertion size was a key factor affecting retrotransposon accumulation, where large elements accumulated in gene poor regions where they were less likely to perturb gene regulation; and 3) insertion patterns surrounding regulatory elements were persistent at the gene level. From this we propose a significant change to the current retrotransposon accumulation model; rather than random insertion constrained by local sequence composition, we propose that insertion is instead primarily constrained by local chromatin structure. Therefore, L1s and SINEs both preferentially insert into gene/regulatory element rich euchromatic domains, where L1s with their relatively high mutational burden are quickly eliminated via purifying selection at a much higher rate than SINEs. Over time this results in an enrichment of SINEs in euchromatic domains and an enrichment of L1s in heterochromatic domains.

## Conclusion

In conjunction with large scale conservation of synteny (Chowdhary et al. 1998), gene regulation (Chan et al. 2009) and the structure of RDs/TADs (Dixon et al. 2012; Ryba et al. 2010),our findings suggest that large scale positional conservation of old and new non-LTR retrotransposons results from their association with the regulatory activity of large genomic domains. Therefore we propose that similar constraints on insertion and accumulation of clade specific retrotransposons in different species can define common trajectories for genome evolution.

## Additional Files

### Additional file 1 — Supplementary information

Figures S1–S31, Tables S1–S6.

### Author’s contributions

R.M.B., R.D.K., J.M.R., and D.L.A. designed research; R.M.B. performed research; and R.M.B., R.D.K., and D.L.A. wrote the paper.

## Acknowledgements

For reviewing our manuscript and providing helpful advice we would like to thank the following: Simon Baxter, Atma Ivancevic and Lu Zeng from the University of Adelaide; Kirsty Kitto from Queensland University of Technology; and Udaya DeSilva from Oklahoma State University.

## Availability of data and materials

All data was obtained from publicly available repositories, urls can be found in supporting material (Table S1–S4). R scripts used to perform analyses can be found at https://github.com/AdelaideBioinfo/retrotransposonAccumulation.

